# Thermostability and binding properties of single-chained Fv fragments derived from therapeutic antibodies

**DOI:** 10.1101/2024.02.09.577534

**Authors:** Takashi Tadokoro, Harumi Tsuboi, Kota Nakamura, Tetsushi Hayakawa, Reo Ohmura, Izumi Kato, Masaki Inoue, Shin-ichi Tsunoda, Sayaka Niizuma, Yukari Okada, Satoko Otsuguro, Katsumi Maenaka

**Affiliations:** Laboratory of Biomolecular Science, Faculty of Pharmaceutical Sciences, Hokkaido University, Sapporo, Japan; Center for Research and Education on Drug Discovery, Faculty of Pharmaceutical Sciences, Hokkaido University, Sapporo, Japan; Laboratory of Cellular and Molecular Physiology, The Faculty of Pharmaceutical Sciences, Kobe Gakuin University, Kobe, Japan; Institute for Vaccine Research and Development (HU-IVReD), Hokkaido University; Sapporo, Japan; Global Station for Biosurfaces and Drug Discovery, Hokkaido University, Sapporo, Japan

**Author notes:** To whom correspondence should be addressed: Katsumi Maenaka: Faculty of Pharmaceutical Sciences, Hokkaido University, Kita-12, Nishi-6, Kita-ku, Sapporo 060-0812, Japan.

**Keywords:** antibody, single chain fragment variable, thermal stability

## Abstract

Small antibody fragments have recently been used as alternatives to full-length monoclonal antibodies in therapeutic applications. One of the most popular fragment antibodies is single-chain fragment variables (scFvs), consisting of variable heavy (V_H_) and variable light (V_L_) domains linked by a flexible peptide linker. scFvs have small molecular sizes, which enables good tissue penetration and low immunogenicity. Despite these advantages, the use of scFvs, especially for therapeutic purpose, is still limited because of the difficulty to regulate the binding activity and conformational stability. In this study, we constructed and analyzed 10 scFv fragments derived from 10 representatives of FDA-approved mAbs to evaluate their physicochemical properties. Differential scanning calorimetry analysis showed that scFvs exhibited relatively high but varied thermostability, from 50 to 70 °C of melting temperatures, and different unfolding cooperativity. Surface plasmon resonance analysis revealed that scFvs fragments that exhibit high stability and cooperative unfolding likely tend to maintain antigen binding. This study demonstrated the comprehensive physicochemical properties of scFvs derived from FDA-approved antibodies, providing insights into antibody design and development.

## Introduction

During the last three decades, monoclonal antibodies (mAbs) have made a dramatic ascend from an experimental therapeutic tool to the most vigorously researched area in the pharmaceutical market (2; 28; 17). One of the drawbacks of mAbs, however, is that full-length antibodies are large (∼150 kDa) and highly complex molecules, which have disadvantages for tissue penetration, production and manipulation. Therefore, recent trend is actively moving towards developing smaller recombinant antibody fragments, as a useful alternative to full-length monoclonal antibodies (9; 22). Among these recombinant antibodies, single-chain variable fragments (scFvs) are the most popular type. This is because they can easily be manipulated using molecular genetic tools. scFvs can be conjugated or produced as bi-specific or multivalent forms and can be produced in *E. coli* expression system (33).

To understand physicochemical properties of mAbs which influence their stability and binding activity is remarkably important for designing highly stable antibodies with excellent affinity and specificity (24; 18; 25). Full-length antibodies comprise a fragment antigen binding (Fab) and a fragment crystallizable (Fc) segment, resulting in V_H_, V_L_, C_L_, C_H_1, C_H_2 and C_H_3 domains. The stability of intact antibodies depends on those of the component domains as well as those of their complexes with the respective interaction interfaces. In contrast to full-length antibodies, scFvs consist only of a variable heavy (V_H_) and variable light (V_L_) domains connected by a flexible linker. Essentially, a scFv maintains a complete antigen-binding site for an antibody, but with the omission of Fc parts. Many applications of scFv fragments under various situations may essentially benefit from an increase in the stability of fragments (34), however, comprehensive set of experimental data of scFvs was limited. Further investigation of the thermostability, ligand-binding affinity of scFvs and its determining factors are needed for the rational design and development of antibody derivatives for the practical use.

In the present study, we selected, constructed and analyzed 10 scFv fragments corresponding to the antigen-binding domains of 10 FDA-approved mAbs, because these mAbs are assumed to possess optimal thermostability and binding properties. Surprisingly, the thermostability of scFvs varied although the thermal denaturation (*T*_m_) range was relatively high, approximately 50–70 °C of melting temperatures (*T*_m_s). Our findings summarize the comprehensive thermostability and binding properties of a diverse set of scFvs and compare them to their parental FDA-approved antibodies, which provides insight into the rational design of fragmented antibodies.

## Results

### Initial analysis of VH and VL fragments and comparison to full-length antibodies

V_H_ and V_L_ domains exhibit mutual stabilization due to their interaction, which depends on the intrinsic stability of each domain as well as the strength of their interface forces (35; 26). Many groups including Buhler *et al.,* as well as our group, have previously reported that the domain orientation of scFv antibody had an impact on its biological and physicochemical properties, and production yield (1; 27). Thus, we first investigated one of the most commonly used FDA-approved mAbs, trastuzumab (Herceptin) (8; 14), for expression yield, thermal stability and antigen binding affinity of its V_H_-linker-V_L_ (HL) and V_L_-linker-V_H_ (LH) scFv forms. To this end, we designed HL-scFv and LH-scFv constructs (**Fig. 1A**), expressed in *E. coli* as inclusion bodies, refolded, and purified using gel filtration chromatography. The chromatograms and the SDS-PAGE are shown in **Fig. 1B and 1C**, respectively, indicating that the resultant proteins were highly pure. The LH construct yielded 11 mg from 1 L culture, whereas the HL one did 2.2 mg, indicating that the LH exhibited roughly 5 times higher yield.

**Figure 1.**
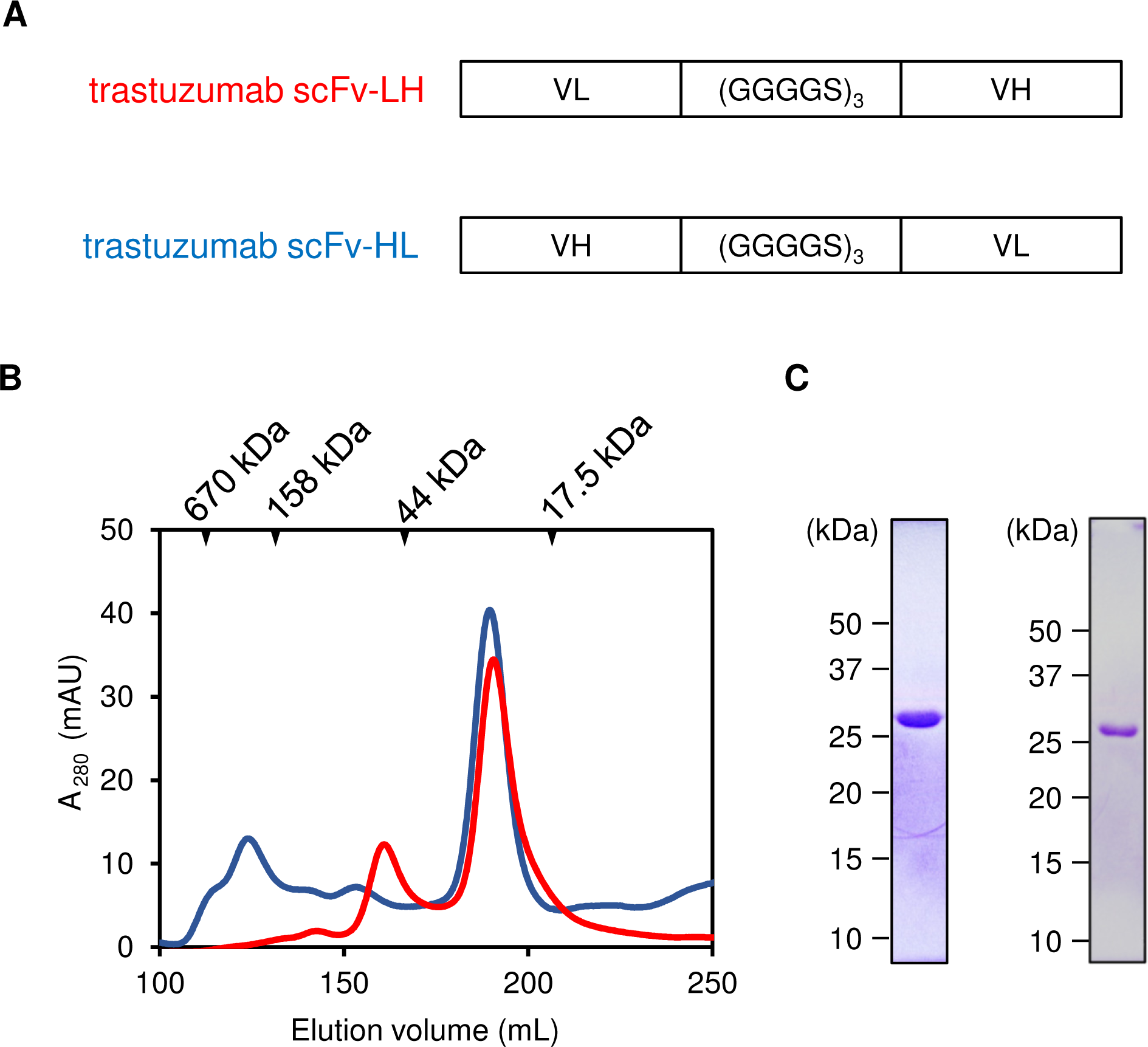
Construction and production of trastuzumab scFvs with V_H_ and V_L_ orientations. A. Schematic representation of the V_H_ and V_L_ constructs including the linker used in this study. B. Chromatograms of the size exclusion chromatography for HL- and LH-scFvs. C. SDS-PAGE of the purified HL- and LH-constructs.

We sought to examine the binding affinities of LH and HL constructs to their target antigen, the extracellular domain of the HER2 receptor (HER2-ECD) (14). Binding analysis using surface plasmon resonance (SPR) showed that both LH and HL constructs exhibited comparable affinity to HER2-ECD, with a K_D_ value of ∼ 0.2 nM, with similar kinetic parameters (**Fig. 2A and B**, **Table 1**).

**Figure 2.**
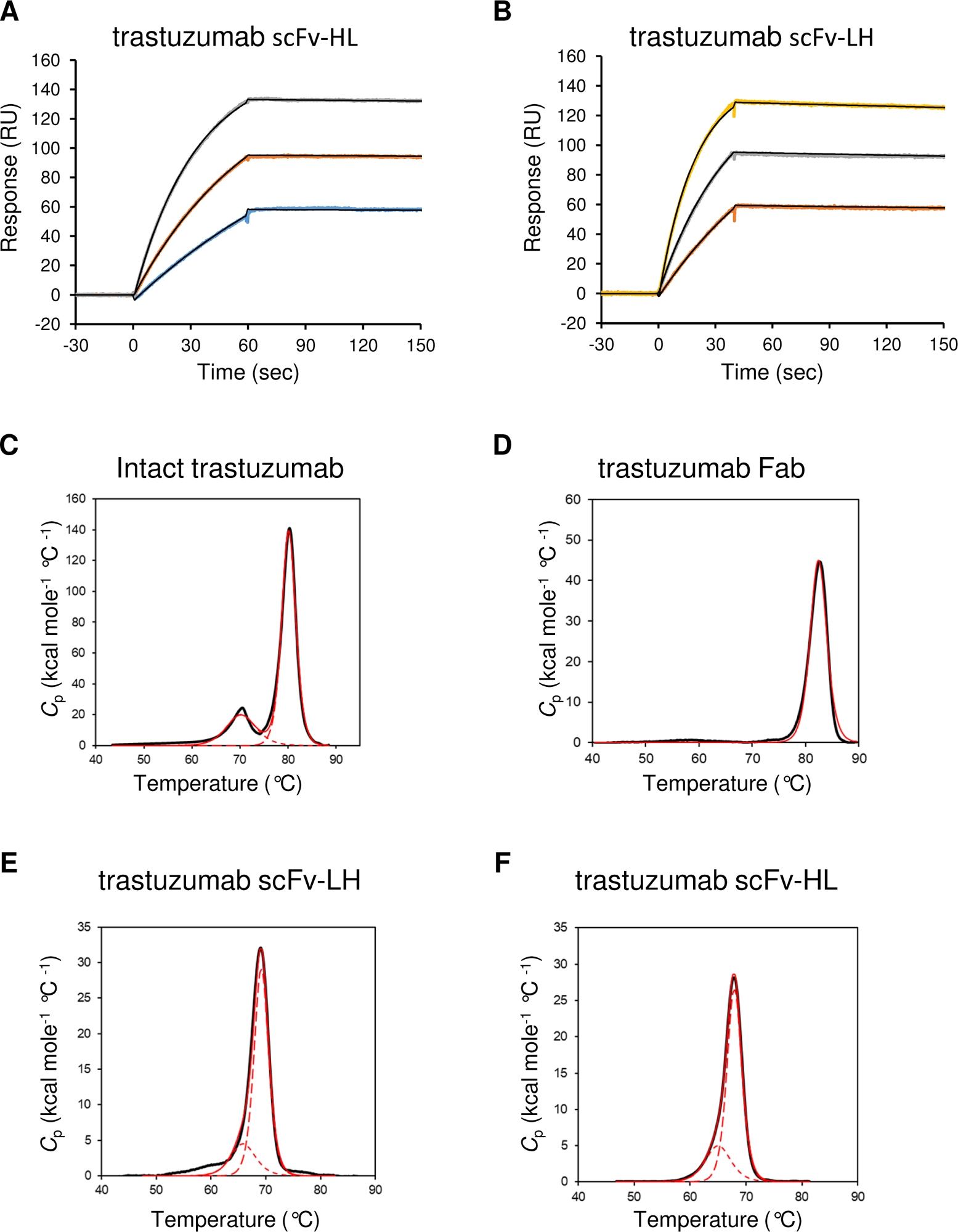
Representative sensorgrams of the binding of trastuzumab scFv to HER2 and differential scanning calorimetry analysis of trastuzumab and its fragments. A. Sensorgrams for scFv-HL. B. Sensorgrams for the scFv-LH. Representative thermograms of C. Full-length trastuzumab, D. Fab fragment of trastuzumab, C. trastuzumab scFv-LH and D. trastuzumab scFv-HL. Experimental curves of intact trastuzumab and its derivatives are indicated with black solid line, whereas the fitting curves of deconvolution results are indicated with red solid line for entire fitting and red broken line for each transition.

**Table 1:** Kinetic parameters of the trastuzumab scFv constructs for HER2 binding.

| | $K_D$ (nM) | $k_{on}$ ($M^{-1} s^{-1}$) | $k_{off}$ ($s^{-1}$) |
| --- | --- | --- | --- |
| scFv-HL | 0.22 | $3.1 \times 10^5$ | $6.7 \times 10^{-4}$ |
| scFv-LH | 0.16 | $3.2 \times 10^5$ | $5.1 \times 10^{-4}$ |

The kinetic parameters of binding were calculated by the fitting with the simple 1:1 Langmuir binding model. The equilibrium dissociation constant, $K_D$ values were calculated with the equation $K_D = k_{off}/k_{on}$ .

Next, using differential scanning calorimetry (DSC), we compared the thermal stability of scFv-HL and scFv-LH fragments with the parental full-length IgG and Fab fragment of trastuzumab. The thermograms were analyzed with non-two-state model, showing that full-length trastuzumab exhibited two peaks, a small peak (*T*_m_^1^ = 72 °C) and a large major peak (*T*_m_^2^ = 82 °C) (**Fig. 2C and Table 2**), which is consistent with previous reports (12; 32). As expected, the trastuzumab Fab fragment showed a single peak at 82 °C (**Fig. 2D**), which corresponded well with the 82 °C major peak on the full-length trastuzumab thermogram. In contrast, the LH and HL constructs both unfolded in a similar manner, with a single peak at *T*_m_ of 69 °C and 68 °C, respectively (**Fig. 2E and F**), suggesting that the stabilities of LH and HL were almost identical.

**Table 2:** Summary of the *T*_m_ values of trastuzumab and derivatives.

| | $T_m^1$ (°C) | $T_m^2$ (°C) | No. of peak |
| --- | --- | --- | --- |
| Full length | 71.8 | 80.1 | multiple |
| Fab | 82.4 | — | single |
| scFv-HL | 65.0 | 67.9 | single |
| scFv-LH | 65.7 | 69.9 | single |
| Experimental DSC curves were fitted with the non-two-state model. |  |  |  |

Taken together, the biophysical characteristics, target binding affinity and conformational stability were similar in both the LH and HL constructs for trastuzumab. On the other hand, since the yield was higher for the LH construct, we decided to use the LH scFvs in all further experiments.

### Construction of scFvs of 10 FDA-approved antibodies

We designed and produced scFvs of ten FDA-approved antibodies, as listed in **Table 3**. The design and production method were essentially the same as for trastuzumab scFv, and the yield for each construct ranged from 0.1 to 10 mg of 1 L culture.

**Table 3:** Kinetic parameters of scFv for antigen binding.

| antibody | Antigen | $K_D$ (nM) | $k_{on}$ ($M^{-1} s^{-1}$) | $k_{off}$ ($s^{-1}$) |
| --- | --- | --- | --- | --- |
| trastuzumab scFv | HER2 | 0.33 | $8.13 \times 10^5$ | $2.71 \times 10^{-4}$ |
| rituximab scFv | CD20 | 149 | $6.94 \times 10^3$ | $1.03 \times 10^{-3}$ |
| cetuximab scFv | EGFR | 1.13 | $2.78 \times 10^5$ | $3.15 \times 10^{-3}$ |
| infliximab scFv | TNFα | 1.75 | $1.04 \times 10^6$ | $1.82 \times 10^{-3}$ |
| adalimumab scFv | TNFα | 1.18 | $7.97 \times 10^5$ | $9.39 \times 10^{-4}$ |
| alemtuzumab scFv | CD52 | 717 | $9.75 \times 10^3$ | $6.99 \times 10^{-3}$ |
| pertuzumab scFv | HER2 | 9.19 | $1.20 \times 10^5$ | $1.10 \times 10^{-3}$ |
| obinutuzumab scFv | CD20 | 1120 | $1.60 \times 10^4$ | $2.72 \times 10^{-2}$ |
| ofatumumab scFv | CD20 | ND | ND | ND |

The kinetic parameters of binding were calculated by the fitting with the simple 1:1 Langmuir binding model. The equilibrium dissociation constant, $K_D$ values were calculated with the equation $K_D = k_{off}/k_{on}$ . ND; Not determined.

We performed surface plasmon resonance (SPR) binding analyses using the following antigens: EGFR for cetuximab and panitumumab, HER2-ECD for trastuzumab and pertuzumab, CD52 for alemtuzumab, CD20 for rituximab, obinutuzumab and ofatumumab, and TNFα for infliximab and adalimumab. The SPR sensorgrams are shown in **Fig. 3**, and the kinetic parameters determined by SPR were summarized in **Table 3**. All the scFvs whose antigen binding affinity was examined retained affinity to their respective antigens, roughly nM to µM level of dissociation constant, *K*_D_ (**Fig. 3**, **Table 3**), besides the ofatumumab scFv. Only ofatumumab scFv did not show an increase in response in a dose-dependent manner (**Fig. 3H**). Among the remaining scFvs, six scFvs showed slow dissociation characteristics, while the fast dissociation kinetics (∼10^−2^ s^−1^) for obinutuzumab scFv resulted in a micromolar *K*_D_ value. These scFvs also showed binding activities comparable to those of intact antibodies (**Supplementary Table 1**).

**Figure 3.**
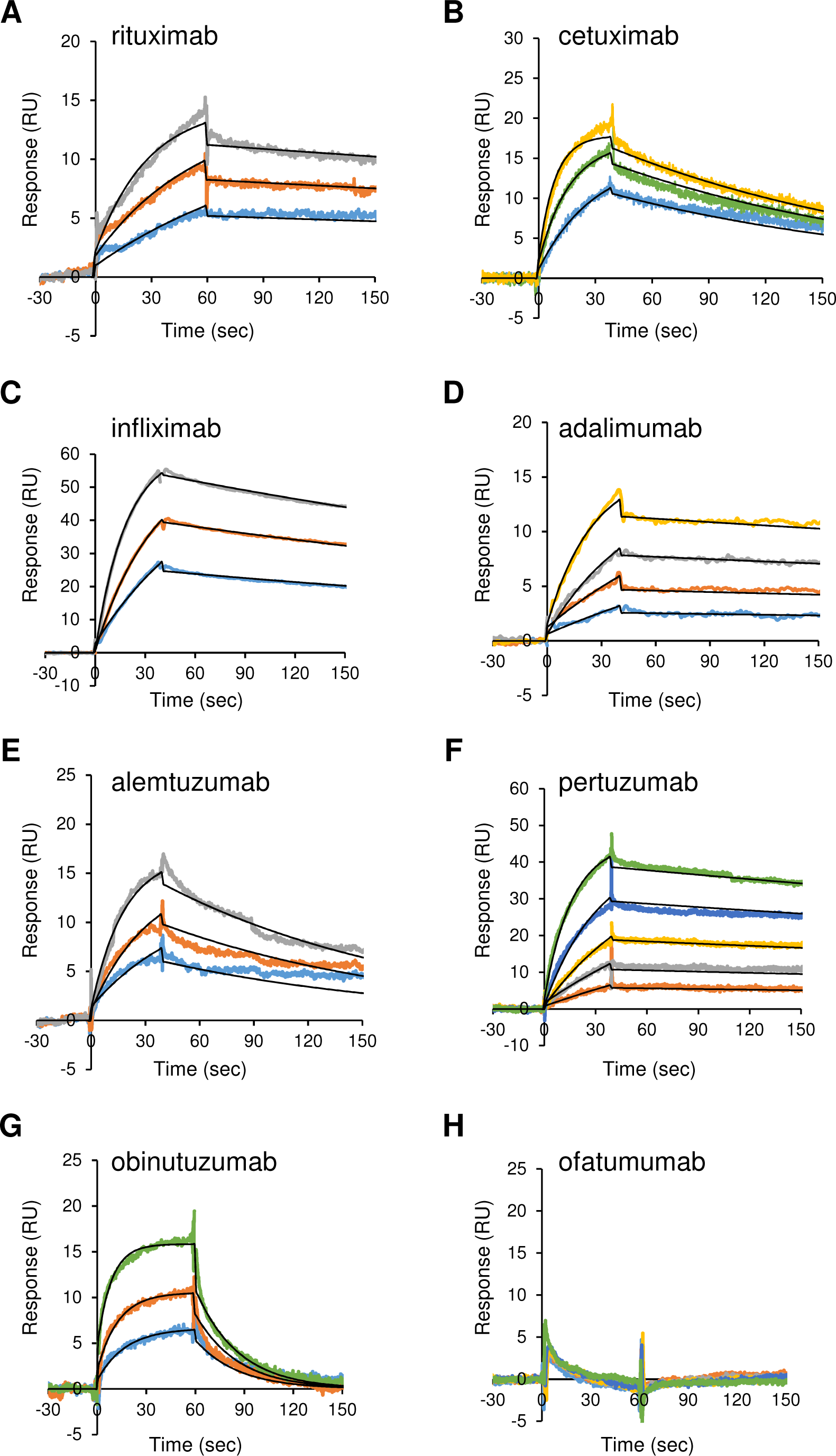
SPR binding analysis of the FDA-approved antibody scFvs to their antigens. Representative sensorgrams are shown. The antibody and antigen pairs are as follows: A; rituximab vs CD20, B; cetuximab vs EGFR, C; infliximab vs TNFα D; adalimumab vs TNFα, E; alemtuzumab vs CD52, F; pertuzumab vs HER2, G; obinutuzumab vs CD20.

### DSC analysis of the scFv fragments

Next, we performed differential scanning calorimetry (DSC) analysis of the 10 scFv fragments. The thermograms were analyzed with the non-two-state model assuming at least two transitions (see **Supplementary Fig. 1**), as none of them could be analyzed with a simple two-state model. The results showed significantly different thermostability profiles among the scFvs, which prompted us to divide the fragments into two groups (**Fig. 4**). Five scFvs (cetuximab, infliximab, mogamulizumab, obinutuzumab, ofatumumab) showed a single clear peak with two partly or mostly overlapping transitions (**Fig. 4B**), whereas the other scFvs (rituximab, adalimumab, alemtuzumab, pertuzumab) showed multiple peaks with two relatively independent transitions (**Fig. 4B**). To further characterize the stability profile, we compared the differences between the two transition temperaturesΔ*T*_m,_ in this study. The Δ*T*_m_ values for a former group were 2.7 °C to 3.3 °C, while those for a latter group were 6.1 to 16.7 °C (**Table 4**). The results suggest that there may be a distinct decrease of cooperativity of the unfolding process when the difference between *T*_m_^1^ and *T*_m_^2^ is roughly more than 5-6 °C.

**Figure 4.**
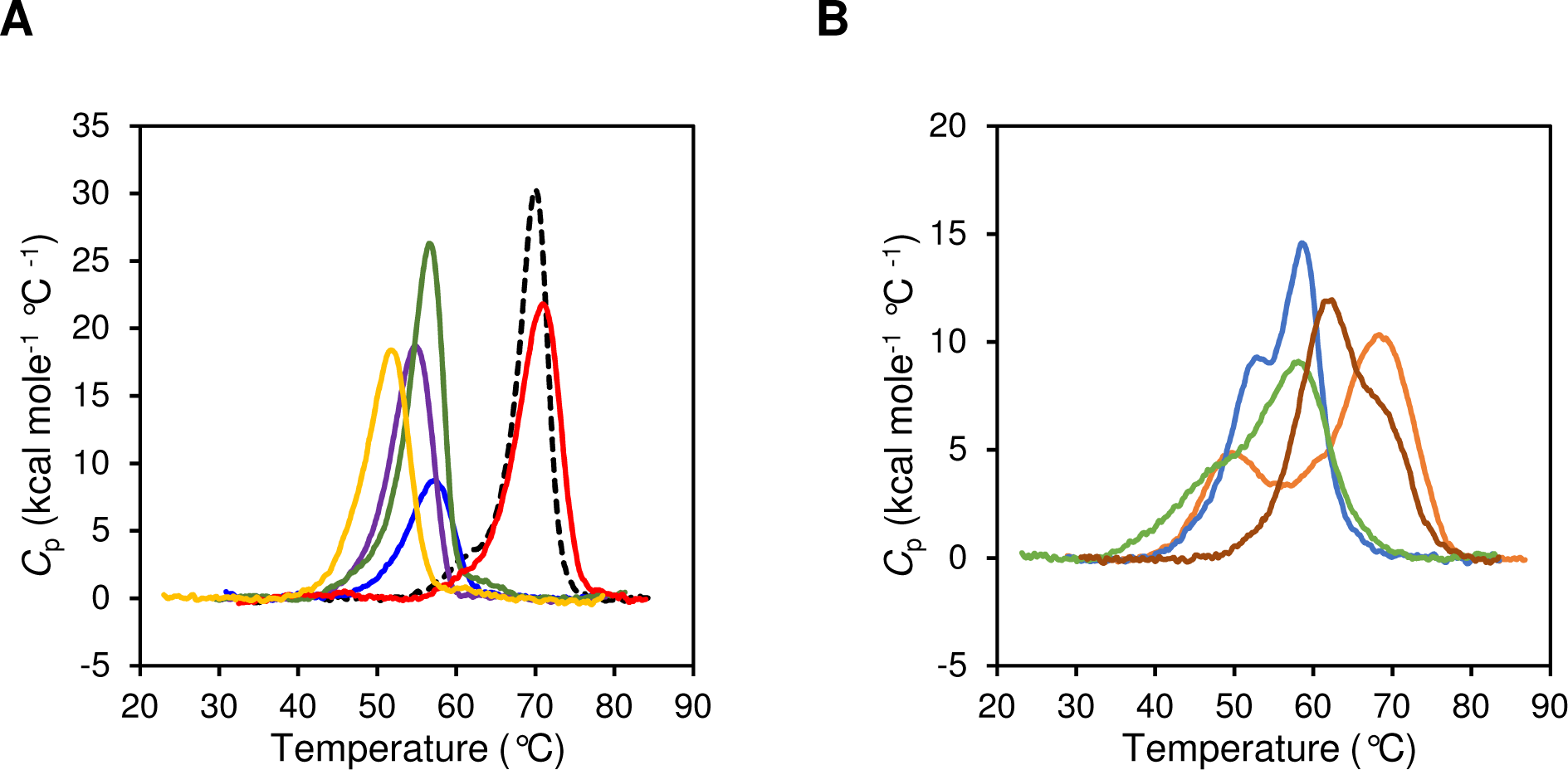
Differential scanning calorimetry analysis of 10 scFv fragments. A. The first group of the fragments with a single sharp peak includes cetuximab (blue), infliximab (purple), mogamulizumab (green), obinutuzumab (red), and ofatumumab (yellow). Trastuzumab (black broken) is also included in this group. B. The second group of the fragments with blunted peaks includes rituximab (orange), adalimumab (cyan), alemtuzumab (light green), pertuzumab (brown).

**Table 4:** Summary of the *T*_m_ values of scFv.

| | $T_m^1$ (°C) | $T_m^2$ (°C) | $\Delta T_m$ (°C) | No. of observed peak |
| --- | --- | --- | --- | --- |
| trastuzumab scFv | 65.7 | 69.9 | 4.2 | single |
| rituximab scFv | 51.3 | 68.0 | 16.7 | multiple |
| cetuximab scFv | 54.7 | 57.7 | 3.0 | single |
| infliximab scFv | 52.4 | 55.3 | 2.9 | single |
| adalimumab scFv | 52.9 | 59.0 | 6.1 | multiple |
| alemtuzumab scFv | 48.9 | 58.4 | 9.5 | multiple |
| mogamulizumab scFv | 53.9 | 56.7 | 2.8 | single |
| pertuzumab scFv | 62.0 | 69.7 | 7.7 | multiple |
| obinutuzumab scFv | 67.9 | 71.2 | 3.3 | single |
| ofatumumab scFv | 49.6 | 52.3 | 2.7 | single |

Experimental DSC curves were analyzed with the non-two-state model assuming two transition processes. The first and second transition temperatures were indicated as $T_m^1$ and $T_m^2$ . We compared the differences between $T_m^1$ and $T_m^2$ ( $\Delta T_m = T_m^2 - T_m^1$ ).

## Discussion

The stabilization of antibodies has been of great interest as one of the specific aspects of antibody engineering over the past few decades. Researchers put several efforts with different strategies, and the approaches can be classified as (a) knowledge-based, (b) statistical, and (c) structure-based methods (18; 28; 34). However, minimization of antibody-related molecules is required. scFvs and VHHs are expected candidates. To understand the factors influencing scFv stability in more detail, followed by developing appropriate rational design methods, further physicochemical understanding of protein stability and function, and correlation of structural and functional properties of scFv are still needed. In this study, we focused on the FDA-approved antibody drugs and compared thermal stability and antigen-binding activity of their scFv fragments. Since FDA-approved antibodies can be assumed to be optimized in terms of parameters necessary for clinical use, including thermostability, one could expect their parts (such as Fabs or scFvs) to exhibit an optimal unfolding profile, akin to trastuzumab Fab. The overall stability of scFv fragments is not thought to show a simple two-state transition in the denaturing process, but is thought to be due to the intrinsic stability of the V_H_ and V_L_ domains, as well as to the stability of the interdomain (34). Thus, we analyzed the DSC data with a non-two-state model. Unexpectedly, we found that approximately half of the antibodies tested in this study display DSC curves with multiple peaks, probably due to the large differences in the stability between V_H_ and V_L_ fragments.

According to our SPR analysis, most scFvs showed nM to µM *K*_D_ levels against their antigens. To compare our results with published data, we summarized the reported kinetic parameters for antigen binding of FDA-approved antibodies examined in this study (**Supplementary Table 1**). Our results showed that rituximab, alemtuzumab and obinutuzumab scFvs exhibited relatively weak binding activity, µM to sub µM of *K*_D_ values, and former two scFvs showed multiple peaks in DSC analysis. As shown in **Supplementary Table 1**, most of the scFvs in this study exhibited comparable dissociation constants, *K*_D_s, for antigen binding with intact or Fab antibodies, besides obinutuzumab although the binding experiment was not SPR analysis. The result of obinutuzumab showed a higher affinity than our result of scFv, probably because of the avidity effect of intact IgG1 (20). In addition, infliximab scFv showed weaker binding activity to its intact IgG form, although the binding affinity (K_D_ = 1.75 nM) remained strong enough. Because the ligand, TNF, is a trimeric protein, likely exhibiting avidity effect for its IgG.

Since a scFv is thought to start losing its function with the first transition in a series of multiple transitions during denaturation, it has been proposed to find and improve stability-limiting properties (35). It is necessary to consider separately in case that one domain is intrinsically stable than the other and that interdomain stability is largely different from individual domain stabilities. Antigen-binding activity may be lost in former case because either of the variable domain denature, whereas maintenance of folded variable domains may allow to interact with antigen regardless of the interface association in the latter case. Therefore, to maintain function, it is presumably important for scFv to maintain stability, such that it is observed as the temperature at the beginning of thermal denaturation and the temperature of the peak of the first transition. In this regard, ofatumumab scFv likely exhibited the lowest temperatures at the beginning and peak of first transition (**Fig. 4**). This may be one of the significant factors that ofatumumab scFv, resulting in the loss of binding activity.

The Fab of trastuzumab exhibited a single peak thermogram with a *T*_m_ of 82 °C, which is almost identical to the *T*_m_ of the major peak in intact trastuzumab. It is generally accepted that the denaturing of Fabs in full-length antibodies and Fab fragments occurs with a single transition. Thus, a comparison of thermostability of scFv and its Fab fragment or Fab domain in the intact form would be helpful to discuss about stability factors although it may be limited to fragments that exhibit single-peak thermograms. Assuming that the *T*_m_^2^ likely corresponds to the peak top of the thermograms as shown in **Fig. 2E** and **2F**, the *T*_m_^2^ for scFvs of trastuzumab is relatively high, approximately 70 °C, as a mammalian globular protein, which differs by about 10 °C lower than that for its Fab form or that for the original Fab domain in the intact form. This is reasonable, because scFv loses one of the two interdomain interactions and interchain disulfide bonds in C_H_1, which effectively stabilizes the Fab fragment. In contrast, the V_H_-V_L_ interaction was stronger, and the thermostability and cooperativity were higher, which presumably contributed to the maintenance of binding activity. Similar to a general protein-protein interaction model, antibody-antigen binding is considered to follow either lock and key model and induced-fit model. For the induced-fit model, a little low cooperativity may be allowed only when the variable domains remain their fold enough to recognize antigen upon losing interdomain stability. According to the previous DSC analyses using intact IgGs, *T*_m_ values derived from Fab domain for intact rituximab, adalimumab and pertuzumab are available to be 76 °C, 70 °C and 77 °C, respectively (15; 29; 23). Of the three antibodies, only pertuzumab scFv showed a single peak transition in DSC analysis, and the *T*_m_^2^ is 69.7 °C, which differs by approximately 7 °C. Rituximab and adalimumab scFvs showed multiple peaks in DSC curves, but the Δ*T*_m_ of adalimumab is relatively small and thus, comparing *T*_m_^2^, 59.0 °C, the difference with the *T*_m_ of Fab is 11 °C. On the other hand, rituximab showed a large Δ*T*_m_ of 16.7 °C, which may not be worth comparing in this way. We have previously shown that the measles neutralizing antibody 2F4 scFv is less stable at approximately 25 °C in *T*_m_ than its Fab, which resulted in approximately 300 times lower binding affinity of scFv than that of its Fab fragment (14).Taken together, loss of the stability derived from C_H_1 domain may be allowed to some extent, around 10 °C in *T*_m_, upon developing scFv fragments.

DSC analysis of intact trastuzumab showed two distinct peaks in the thermogram, with *T*_m_ values of 72 °C and 82 °C. IgG1 antibody typically has three peaks in DSC measurement, *T*_m_^1^, *T*_m_^2^ and *T*_m_^3^ presumably corresponding to the unfolding of C_H_2 domain, Fab regions, and C_H_3 domain, respectively (7). Thus, thermal unfolding of the C_H_3 domain may occur simultaneously when the Fab domain of trastuzumab unfolds. Vermeer *et al*. studied the unfolding of the present IgG of isotype 2b, mainly by DSC and circular dichroism (CD) spectroscopy (30; 31). They showed that the unfolding of IgG is a complex process, and is characterized by the sum effect of mainly two transitions of Fab and Fc fragments and several, at least partly independent, intermediated state.

To facilitate manufacturing and storage, as well as to promote long serum half-life, expanding the range of applications for practical use, biophysical properties such as thermostability are often limited by antibody variable domains, that differ greatly in their intrinsic properties and are responsible for antigen specificity. Our study provides insights into the rational design of stable antibody fragments suitable for clinical use.

## Materials and Methods

### Preparation of trastuzumab Fab

Trastuzumab was purchased from Roche. Trastuzumab Fab was prepared using Pierce Fab Preparation Kit (Thermo Fisher Scientific). The digestion and purification were performed following the manufacturer’s instruction. Briefly, 2 mg of intact trastuzumab was applied to a spin column containing papain-immobilized resin that was pre-equilibrated with cysteine-containing digestion buffer. For the digestion reaction, the column was incubated at 37 °C for 4 h. The digestion product was separated from papain by centrifugation. Subsequently, the digest was applied to a PBS-equilibrated Protein A-immobilized agarose column to capture the Fc fragment and undigested full-length trastuzumab. The Fab fragment was collected as a flow-through fraction.

### Recombinant scFv preparation

DNA encoding 11 scFv fragments was chemically synthesized by Eurofins Scientific and was inserted into NdeI-HindIII site of pET22b vector. The resulting expression plasmids were introduced into the *E. coli* BL21(DE3)pLysS strain. The transformants were cultured in 2×YT medium 100 mg/L ampicillin (Nacalai Tesque) at 37 °C until the OD_600_ reached 0.4-0.6, and isopropyl β-d-thiogalactopyranoside (IPTG) (Nacalai Tesque) was then added at a final concentration of 1 mM. The bacteria were further cultivated for 4 h, and harvested by centrifugation at 5,000×g for 10 min. The pellet was resuspended in buffer (50 mM TrisHCl pH8.0, 100 mM NaCl), sonicated, and the resulting scFvs were washed repeatedly with Triton wash buffer (50 mM Tris-HCl pH 8.0, 100 mM NaCl, 0.5% Triton X-100). Purified inclusion bodies were solubilized in denaturation buffer (50 mM Tris-HCl pH 8.0, 10 mM EDTA, 6 M guanidine HCl) and slowly diluted with the ice-cold refolding buffer (100 mM Tris-HCl, pH 8.0, 2 mM ethylenediaminetetraacetic acid (EDTA), 0.4 M L-Arginine, 3.73 mM cystamine, 6.37 mM cystamine) to a final protein concentration of 1.5 μM. After incubation for 72 h at 4 °C, the refolded protein was concentrated using a VIVAFLOW50 system (Sartorius). The concentrated sample was filtrated with 0.22 µm membrane filter. The soluble fraction was separated by centrifugation at 30,000×g for 30 min and subjected to size exclusion chromatography using a HiLoad26/60 Superdex75 column (GE Healthcare) with gel filtration buffer (20 mM Tris-HCl pH 8.0, 100 mM NaCl). Fractions containing scFv protein were pooled and concentrated to ∼1.0 mg/mL. Protein concentration was determined using UV spectrophotometer and the purity was checked by SDS-PAGE.

### Preparation of the antigen proteins

EGFR, which is the antigen for cetuximab and panitumumab, were prepared with baculobvirus-silkworm expression system according to the previous report (6). A human erbB1 cDNA fragment encoding residues 1–618 of the mature sequence (extracellular domain (ECD) of EGFR), followed by a hexahistidine tag and stop codon, was subcloned into pFastBac1 purchased from Invitrogen and designated as pFastBac1-EGFR-ECD. *E. coli* BmDH10Bac was previously constructed in our laboratory (21). *E. coli* BmDH10Bac were transformed with pFastBac1-EGFR-ECD. Positive transformants were chosen by blue-white selection and colony PCR. Appropriate colonies were grown and the recombinant bacmid DNA was extracted using the NucleoBond Xtra Midi plasmid purification kit (TAKARA). The fifth instar silkworm larvae, purchased from Ehime-Sanshu (Japan), were grown at 25 °C with moderate humidity in an incubator. Recombinant bacmid DNA was diluted in 50 µ L of ultra-pure water, and mixed with 3 µ L of DMRIE-C (Invitrogen). After incubation at room temperature for 45 min, the bacmid-DMRIE-C mixture was injected into the fifth instar silkworm larvae. Six days after the bacmid injection, silkworm hemolymph was collected and immediately mixed with sodium thiosulfate at final concentration of 0.5%. The hemolymph mixture was subsequently centrifuged at 10,000 x g for 10 min to remove hemolymph debris and filtered through a 0.45 µm PVDF membrane filter (Millipore). The filtered fraction was centrifuged at 45,000× g for 30 min.

The resultant supernatant, containing the target protein, was precipitated at 80% saturation with ammonium sulfate to remove lipids and debris. The precipitates were collected by centrifugation at 20,000 ×g for 30 min, solubilized in Tris buffered saline (TBS) containing 25 mM imidazole (20 mM Tris-HCl pH 8.0, 150 mM NaCl, 25 mM imidazole), and applied to Ni-NTA column (GE Healthcare) equilibrated with the same buffer. The protein was eluted from the column using elution buffer containing 300 mM imidazole. Fractions containing the target proteins were pooled, concentrated and applied to a Superose6 GL gel filtration column (GE Healthcare) equilibrated with gel filtration buffer (20 mM Tris-HCl pH 8.0, 150 mM NaCl). Fractions containing pure protein were pooled and used for subsequent analyses.

Recombinant TNFαprotein for infliximab and adalimumab was purified previously (11). Recombinant human HER2-ECD Fc chimera protein for trastuzumab and pertuzumab, and CD52 protein for alemtuzumab were purchased from R&D systems, and recombinant human CD20 for rituximab, obinutuzumab and ofatumumab was purchased from Abcam.

### Binding analysis using surface plasmon resonance

Surface plasmon resonance (SPR) experiments were performed using a BIAcore 3000 (GE Healthcare). The ligand protein was immobilized on a CM5 sensor chip using amine coupling method. For binding analysis, scFv was injected over an immobilized cognate antigen or BSA (control). The binding response at each concentration of scFv was calculated by subtracting the equilibrium response measured in the control-flow cell from the response in each sample-flow cell. The kinetic parameters were calculated by using the curve-fitting facility of the BIAevaluation version 4.1.1 software (GE healthcare) and the simple 1:1 Langmuir binding model (A + B ↔ AB).

### Differential scanning calorimetry (DSC)

The thermal stability of the scFvs was measured using a Capillary VP-DSC system (Malvern) with a cell volume of 0.135 mL. The protein was dissolved in 20 mM sodium phosphate buffer pH 7.0 at 3.4∼6.0 μM. Measurements were performed with a temperature range from 20 °C to 90 °C at a scan rate of 60 °C/hour. Data were analyzed using Origin software version 7.0 (OriginLab).

## Author Contributions

Conceptualization: TT, KM

Resources: KM

Methodology: TT

Investigation and Validation: TT, HT, KN, TH, RO, IK, SN, YO, SO, MI, ST

Visualization: TT

Funding acquisition: KM, TT

Project administration: KM

Supervision: KM

Writing –TT, KM

## Funding

Scientific Research on Innovative Areas and International Group from the MEXT/JSPS KAKENHI [JP20H05873 (K.M)], Japan Agency for Medical Research and Development (AMED) [JP20ae0101047, JP21fk0108463, JP21am0101093, JP22ama121037, JP223fa627005 (K.M)], Takeda Science Foundation (K.M), Hokkaido University, Global Facility Center (GFC), Pharma Science Open Unit (PSOU), funded by MEXT/JST under “Support Program for Implementation of New Equipment Sharing System” (to KM).

Grant-in-Aid for Scientific Research (C) from JSPS KAKENHI [JP 20K05721 (T.T)].

## Conflict of interest

The authors declared no conflict of interest.

## Abbreviations

scFv: single chain fragment variable
mAb: monoclonal antibody
Fab: fragment antigen binding
FDA: U.S. Food and Drug Administration
SPR: Surface plasmon resonance
K_D_: dissociation constant
DSC: differential scanning calorimetry
*T*_m_: melting temperature

## Notes

### Competing Interest Statement

The authors have declared no competing interest.

## References

1. Buhler P, Wetterauer D, Gierschner D, Wetterauer U, Beile U, Wolf P (2010) Influence of Structural Variations on Biological Activity of Anti-PSMA scFv and Immunotoxins Targeting Prostate Cancer. Anticancer Research 30:3373–3379. PMID: WOS:000283009400021 {Medline}

2. Buss N, Henderson S, McFarlane M, Shenton J, de Haan L (2012) Monoclonal antibody therapeutics: history and future. Current Opinion in Pharmacology 12:615–622. PMID: WOS:000310478800016 {Medline}

3. Donaldson J, Kari C, Fragoso R, Rodeck U, Williams J (2009) Design and development of masked therapeutic antibodies to limit off-target effects Application to anti-EGFR antibodies. Cancer Biology & Therapy 8:2147–2152. PMID: WOS:000281477800018 {Medline}

4. Donaldson J, Zer C, Avery K, Bzymek K, Horne D, Williams J (2013) Identification and grafting of a unique peptide-binding site in the Fab framework of monoclonal antibodies. Proceedings of the National Academy of Sciences of the United States of America 110:17456–17461. PMID: WOS:000325943300065 {Medline}

5. Ernst J, Li H, Kim H, Nakamura G, Yansura D, Vandlen R (2005) Isolation and characterization of the B-cell marker CD20. Biochemistry 44:15150–15158. PMID: WOS:000233453000007 {Medline}

6. Ferguson K, Darling P, Mohan M, Macatee T, Lemmon M (2000) Extracellular domains drive homo-but not heterodimerization of erbB receptors. Embo Journal 19:4632–4643. PMID: WOS:000089275600021 {Medline}

7. Garber E, Demarest S (2007) A broad range of Fab stabilities within a host of therapeutic IgGs. Biochemical and Biophysical Research Communications 355:751–757. PMID: WOS:000245001300025 {Medline}

8. Hashiguchi T, Kajikawa M, Maita N, Takeda M, Kuroki K, Sasaki K, Kohda D, Yanagi Y, Maenaka K (2007) Crystal structure of measles virus hemagglutinin provides insight into effective vaccines. Proceedings of the National Academy of Sciences of the United States of America 104:19535–19540. PMID: WOS:000251525800066 {Medline}

9. Holliger P, Hudson P (2005) Engineered antibody fragments and the rise of single domains. Nature Biotechnology 23:1126–1136. PMID: WOS:000231790600030 {Medline}

10. Hu S, Liang S, Guo H, Zhang D, Li H, Wang X, Yang W, Qian W, Hou S, Wang H, Guo Y, Lou Z (2013) Comparison of the Inhibition Mechanisms of Adalimumab and Infliximab in Treating Tumor Necrosis Factor alpha-Associated Diseases from a Molecular View. Journal of Biological Chemistry 288:27059–27067. PMID: WOS:000330597300007 {Medline}

11. Inoue M, Ando D, Kamada H, Taki S, Niiyama M, Mukai Y, Tadokoro T, Maenaka K, Nakayama T, Kado Y, Inoue T, Tsutsumi Y, Tsunoda S (2017) A trimeric structural fusion of an antagonistic tumor necrosis factor-alpha mutant enhances molecular stability and enables facile modification. Journal of Biological Chemistry 292:6438–6451. PMID: WOS:000399813400002 {Medline}

12. Ionescu R, Vlasak J, Price C, Kirchmeier M (2008) Contribution of variable domains to the stability of humanized IgG1 monoclonal antibodies. Journal of Pharmaceutical Sciences 97:1414–1426. PMID: WOS:000254428400007 {Medline}

13. Kaymakcalan Z, Sakorafas P, Bose S, Scesney S, Xiong L, Hanzatian D, Salfeld J, Sasso E (2009) Comparisons of affinities, avidities, and complement activation of adalimumab, infliximab, and etanercept in binding to soluble and membrane tumor necrosis factor. Clinical Immunology 131:308–316. PMID: WOS:000265420000014 {Medline}

14. Khalili H, Godwin A, Choi J, Lever R, Brocchini S (2012) Comparative Binding of Disulfide-Bridged PEG-Fabs. Bioconjugate Chemistry 23:2262–2277. PMID: WOS:000311325000014 {Medline}

15. Lee J, Song C, Kim D, Park I, Lee S, Lee Y, Kim B (2013) Glutamine (Q)-peptide screening for transglutaminase reaction using mRNA display. Biotechnology and Bioengineering 110:353–362. PMID: WOS:000312945800001 {Medline}

16. Liang S, Dai J, Hou S, Su L, Zhang D, Guo H, Hu S, Wang H, Rao Z, Guo Y, Lou Z (2013) Structural Basis for Treating Tumor Necrosis Factor alpha (TNF alpha)-associated Diseases with the Therapeutic Antibody Infliximab. Journal of Biological Chemistry 288:13799–13807. PMID: WOS:000318850300061 {Medline}

17. Lu RM, Hwang YC, Liu IJ, Lee CC, Tsai HZ, Li HJ, Wu HC (2020) Development of therapeutic antibodies for the treatment of diseases. J Biomed Sci 27:1. PMID: 31894001 {Medline}

18. McConnell A, Zhang X, Macomber J, Chau B, Sheffer J, Rahmanian S, Hare E, Spasojevic V, Horlick R, King D, Bowers P (2014) A general approach to antibody thermostabilization. Mabs 6:1274–1282. PMID: WOS:000346878500016 {Medline}

19. Milazzo F, Anastasi A, Chiapparino C, Rosi A, Leoni B, Vesci L, Petronzelli F, De Santis R (2017) AvidinOX-anchored biotinylated trastuzumab and pertuzumab induce down-modulation of ErbB2 and tumor cell death at concentrations order of magnitude lower than not-anchored antibodies. Oncotarget 8:22590–22605. PMID: WOS:000398211100027 {Medline}

20. Mossner E, Brunker P, Moser S, Puntener U, Schmidt C, Herter S, Grau R, Gerdes C, Nopora A, van Puijenbroek E, Ferrara C, Sondermann P, Jager C, Strein P, Fertig G, Friess T, Schull C, Bauer S, Dal Porto J, Del Nagro C, Dabbagh K, Dyer M, Poppema S, Klein C, Umana P (2010) Increasing the efficacy of CD20 antibody therapy through the engineering of a new type II anti-CD20 antibody with enhanced direct and immune effector cell-mediated B-cell cytotoxicity. Blood 115:4393–4402. PMID: WOS:000278372600016 {Medline}

21. Motohashi T, Shimojima T, Fukagawa T, Maenaka K, Park E (2005) Efficient large-scale protein production of larvae and pupae of silkworm by Bombyx mori nuclear polyhedrosis virus bacmid system. Biochemical and Biophysical Research Communications 326:564–569. PMID: WOS:000226143200007 {Medline}

22. Nelson A (2010) Antibody fragments Hope and hype. Mabs 2:77–83. PMID: WOS:000281387700010 {Medline}

23. Noda M, Ishii K, Yamauchi M, Oyama H, Tadokoro T, Maenaka K, Torisu T, Uchiyama S (2019) Identification of IgG1 Aggregation Initiation Region by Hydrogen Deuterium Mass Spectrometry. Journal of Pharmaceutical Sciences 108:2323–2333. PMID: WOS:000477753400012 {Medline}

24. Pantazes RJ, Grisewood MJ, Maranas CD (2011) Recent advances in computational protein design. Curr Opin Struct Biol 21:467–472. PMID: 21600758 {Medline}

25. Rigoldi F, Donini S, Redaelli A, Parisini E, Gautieri A (2018) Review: Engineering of thermostable enzymes for industrial applications. APL Bioeng 2:011501. PMID: 31069285 {Medline}

26. Röthlisberger D, Honegger A, Plückthun A (2005) Domain interactions in the Fab fragment: a comparative evaluation of the single-chain Fv and Fab format engineered with variable domains of different stability. J Mol Biol 347:773–789. PMID: 15769469 {Medline}

27. Tadokoro T, Jahan L, Ito Y, Tahara M, Chen S, Imai A, Sugimura N, Yoshida K, Saito M, Ose T, Hashiguchi T, Takeda M, Fukuhara H, Maneaka K (2020) Biophysical characterization and single-chain Fv construction of a neutralizing antibody to measles virus. Febs Journal 287:145–159. PMID: WOS:000478457100001 {Medline}

28. Tiller K, Tessier P, Yarmush M (2015) Advances in Antibody Design. Annual Review of Biomedical Engineering, Vol 17 17:191–216. PMID: WOS:000367291800008 {Medline}

29. Toughiri R, Wu X, Ruiz D, Huang F, Crissman J, Dickey M, Froning K, Conner E, Cujec T, Demarest S (2016) Comparing domain interactions within antibody Fabs with kappa and lambda light chains. Mabs 8:1276–1285. PMID: WOS:000385569400009 {Medline}

30. Vermeer A, Norde W (2000) The thermal stability of immunoglobulin: Unfolding and aggregation of a multi-domain protein. Biophysical Journal 78:394–404. PMID: WOS:000084772900036 {Medline}

31. Vermeer AW, Norde W, van Amerongen A (2000) The unfolding/denaturation of immunogammaglobulin of isotype 2b and its F(ab) and F(c) fragments. Biophys J 79:2150–2154. PMID: 11023918 {Medline}

32. Wakankar AA, Feeney MB, Rivera J, Chen Y, Kim M, Sharma VK, Wang YJ (2010) Physicochemical stability of the antibody-drug conjugate Trastuzumab-DM1: changes due to modification and conjugation processes. Bioconjug Chem 21:1588–1595. PMID: 20698491 {Medline}

33. Weisser NE, Hall JC (2009) Applications of single-chain variable fragment antibodies in therapeutics and diagnostics. Biotechnol Adv 27:502–520. PMID: 19374944 {Medline}

34. Worn A, Pluckthun A (2001) Stability engineering of antibody single-chain Fv fragments. Journal of Molecular Biology 305:989–1010. PMID: WOS:000166921900001 {Medline}

35. Wörn A, Plückthun A (1998) Mutual stabilization of VL and VH in single-chain antibody fragments, investigated with mutants engineered for stability. Biochemistry 37:13120–13127. PMID: 9748318 {Medline}

