## Supplementary Figure 1 and Supplementary Table 1 for "Thermostability and binding properties of single-chained Fv fragments derived from therapeutic antibodies"

**Supplementary materials**

・Supplementary Figure 1

・Supplementary table 1

**Figure legends**

**Supplementary Fig. 1**

DSC analysis of FDA-approved antibody scFvs related to Fig. 4. Experimental curves of are indicated with black solid line, whereas the fitting curves of deconvolution results are indicated with red solid line for both entire fitting and each transition.

### Supplementary Figure 1

DSC analysis of FDA-approved antibody scFvs related to Fig. 4. Experimental curves of are indicated with black solid line, whereas the fitting curves of deconvolution results are indicated with red solid line for both entire fitting and each transition.

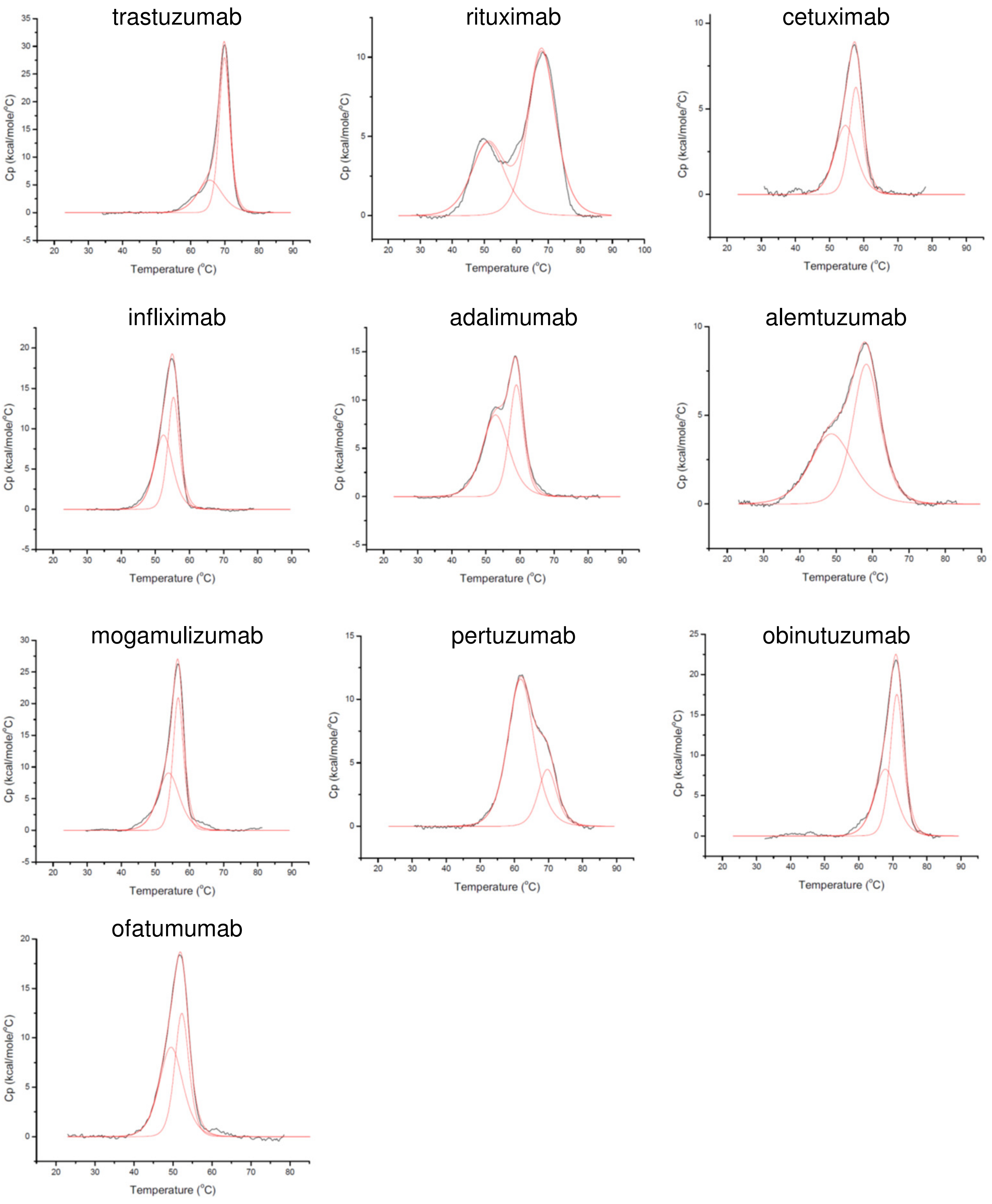

**Supplementary Table 1: Summary of the reported kinetic parameters of full length and fragment antibodies.**

| antibody | | $K_D$ (nM) | $k_{on}$ ($\times10^4$) ($M^{-1}$ $s^{-1}$) | $k_{off}$($\times10^{-5}$) ($s^{-1}$) | ligand / analyte | antigen | Reference |
|---|---|---|---|---|---|---|---|
| trastuzumab | scFv | 0.33 | 81.3 | 27.1 | antigen / antibody | HER2-ECD-Fc | This work |
| | full | 0.11 | 38 | 4.2 | antigen / antibody | human HER2-ECD | (8), (14) |
| | Fab | 0.13 | 102 | 11.5 | antigen / antibody | human HER2-ECD | |
| | scFv | 103.5 | | | antigen / antibody | biotinylated HER2 fragment | |
| rituximab | scFv | 149 | 0.694 | 103 | antigen / antibody | Human CD20 | This work |
| | full | 160 | 0.25 | 41 | antibody / antigen | human CD20 His-tagged protein | (5) |
| | Fab | 280 | 0.17 | 48 | antibody / antigen | human CD20 His-tagged protein | |
| cetuximab | scFv | 1.13 | 27.8 | 315 | antigen / antibody | EGFR ECD | This work |
| | full | 1.9 | | | antigen / antibody | EGFR dIII | (18), (4), (3) |
| | Fab | 0.76 | 260 | 200 | antigen / antibody | EGFR dIII | |
| | scFv | 110 | | | antigen / antibody | EGFR dIII | |
| infliximab | scFv | 1.75 | 104 | 182 | antigen / antibody | human TNFα | This work |
| | full | 0.0273 | 240 | 6.56 | antibody / antigen | soluble TNF | (13), (16) |
| | Fab | 8.70 | 3.50 | 30 | antibody / antigen | human TNFα | |
| adalimumab | scFv | 1.18 | 79.7 | 93.9 | antigen / antibody | human TNFα | This work |
| | full | 0.49 | | | antibody / antigen | human TNFα | (18), (10) |
| | Fab | 0.1152 | 47.84 | 5.512 | antibody / antigen | human TNFα | |
| pertuzumab | scFv | 9.19 | 12.0 | 110 | antigen / antibody | HER2-ECD-Fc | This work |
| | full | 3.8 | 2.86 | 11 | antibody / antigen | HER2-ECD | (19) |
| obinutuzumab | scFv | 1120 | 1.60 | 2720 | antigen / antibody | CD20 | This work |
| | full | 4* | | | | non-Hodgkin lymphoma cells | (20) |

Blank for kinetic parameters were not described in the references.

* $K_D$ value was determined by Scatchard analysis of binding experiments using non-Hodgkin lymphoma cells and labeled antibody with bivalent mode.
